# Three-dimensional in vitro model of the device-tissue interface reveals innate neuroinflammation can be mitigated by antioxidant ceria nanoparticles

**DOI:** 10.1101/2021.11.06.467561

**Authors:** Elaina Atherton, Yue Hu, Sophie Brown, Emily Papiez, Vivian Ling, Vicki L. Colvin, David A. Borton

**Author notes:** Corresponding Author. Address: Brown University, 184 Hope St, Providence, RI 02912, USA Email Address (David A. Borton).

## Abstract

The recording instability of neural implants due to neuroinflammation at the device-tissue interface (DTI) is a primary roadblock to broad adoption of brain-machine interfaces. While a multiphasic immune response, marked by glial scaring, oxidative stress (OS), and neurodegeneration, is well-characterized, the independent contributions of systemic and local “innate” immune responses are not well-understood. Three-dimensional primary neural cultures provide a unique environment for studying the drivers of neuroinflammation by decoupling the innate and systemic immune systems, while conserving an endogenous extracellular matrix and structural and functional network complexity. We created a three-dimensional in vitro model of the DTI by seeding primary cortical cells around microwires. Live imaging of microtissues over time revealed independent innate neuroinflammation, marked by increased OS, decreased neuronal density, and increased functional connectivity. We demonstrated the use of this model for therapeutic screening by directly applying drugs to neural tissue, bypassing low bioavailability through the in vivo blood brain barrier. As there is growing interest in long-acting antioxidant therapies, we tested efficacy of “perpetual” antioxidant ceria nanoparticles, which reduced OS, increased neuronal density, and protected functional connectivity. Overall, our avascular in vitro model of the DTI exhibited symptoms of OS-mediated innate neuroinflammation which were mitigated by antioxidant intervention.

## Introduction

Today, neurotechnology is helping people with paralysis regain function through brain control of prostheses, communication interfaces, and even one’s own limbs^1–4^. Facile control of effectors with single unit activity relies on physical proximity of the device to the local electric fields generated by neurons as they communicate with each other, necessitating probe implantation directly into the brain. Such invasive implantation results in acute tissue damage and chronic inflammation, which eventually disrupts the recording capabilities of the probe over weeks to months.

Inconsistent control of brain-machine interfaces (BMIs) is attributed in part to degradation of recording signal quality, initiated by inflammation at the device-tissue interface^5–7^. Inserting a microelectrode into neural tissue damages microvasculature, breaking the blood brain barrier (BBB) and introducing blood-born immune cells into the area^8^. These systemic immune cells initiate an acute wound healing process through recruitment of microglia, the brain’s local “innate” immune cell^8,9^, resulting in a combined immune reaction at the device-tissue interface marked by a glial scar that encapsulates the damaged tissue and electrode^10–15^. While glial scarring stabilizes at 6 weeks^13^, reactive glia within the scar continue to attack the implanted foreign body through overproduction of reactive oxygen species (ROS) in a process called “frustrated phagocytosis”^16^. This overproduction of ROS causes oxidative stress in the tissues surrounding the implant^17^, inducing neurodegeneration and degradation of implanted materials^10,18–20^. These combined effects contribute to a reduction of recording signal quality and eventual failure of the device^21,22^. Chronic vascular instability caused by repeated micromotion injury^23,24^ and oxidative damage to the BBB^20,25,26^ serves to prolong the acute neuroinflammatory response. As such, chronic neuroinflammation at the device-tissue interface is characterized by a complex interplay between the blood-born systemic immune system and the local “innate” immune system. However, the independent role of the innate immune response in the absence of vascular instability has not been explored.

To better understand the role of the innate immune system in the neuroinflammatory response to recording implants, we engineered an avascular 3D in vitro model of the device-tissue interface to decouple the innate immune response from vascular derived systemic immune drivers. A previously developed two dimensional (2D) in vitro model of the device-tissue interface exhibited histological changes around implanted wires which recapitulated some, but not all, hallmarks of the foreign body response to implants^27,28^. Recent advances in culturing techniques have produced self-assembled three-dimensional (3D) cultures which increase physiological relevance of in vitro paradigms compared to traditional 2D cultures^29–31^. Primary 3D neural cultures produce an endogenous extracellular matrix with multicellular functional and structural networks^30–36^. Such a 3D in vitro paradigm provides easy optical access for live longitudinal imaging of cellular behavior, making it particularly well suited for investigating progressive multicellular processes like neuroinflammation.

Here, we demonstrate the utility of a physiologically relevant 3D in vitro model of the device-tissue interface. The system can be used as a paradigm for studying the mechanisms of immune responses to neural devices, as well as a platform for screening immunomodulating drug treatments. We treated the in vitro device-tissue interface via direct application of an antioxidant to investigate the potential therapeutic efficacy as well as the role of ROS in mediating innate neuroinflammation. In vivo, short-acting antioxidant treatments have been shown to prevent oxidative damage to neural tissue by directly scavenging free radicals. However, long-term stabilization of the device-tissue interface with conventional antioxidants is limited by the temperature instability and high dose requirements of small molecules^20,37–41^. Here, we investigate the use of a long-acting and thermally stable antioxidant made from cerium oxide (Ceria, CeO_2_). This rare earth metal oxide exhibits “perpetual” antioxidant properties in its nanoparticle form^42^, effectively lengthening the time course of treatment at physiological temperatures and decreasing dose requirements. The antioxidant behavior of these materials is derived from the oxygen vacancy clusters present at the cerium(III)-rich nanoparticle surfaces^43^, features that have been shown to be neuroprotective by neutralizing ROS and preventing oxidative damage to neurons^44,45^. Further, ceria nanoparticle treatment has increased neuronal survival in several neuroinflammatory disease states, including Alzheimer’s Disease and traumatic brain injury^46–49^. These previous applications indicate that ceria nanoparticles are capable of mitigating ROS-mediated neuroinflammation and, thus, the addition of ceria to a 3D in vitro model of the device-tissue interface will help elucidate the role of ROS in the independent innate neuroinflammatory response.

We created an in vitro device-tissue interface by seeding cells around a tungsten microwire to model innate neuroinflammation at the exposed recording contact of an implanted electrode. Such an implantation method creates a device-tissue interface without inducing a confounding stab wound. Neuroinflammation was characterized by measuring oxidative stress via live lipid peroxidation, as well as the functional and structural changes in neurons via a genetically encoded calcium indicator with a secondary fluorescent control tag expressed in neurons under the human synapsin1 promoter. Chemical, structural, and functional changes around implants were imaged between 14 and 18 days in vitro (DIV) and compared to unimplanted controls. We found increased lipid peroxidation and decreased neuronal density within 250µm of the microwire implant. Imaging of calcium transients revealed significant increases in functional connectivity and disruption of community structure of neuronal microcircuits in implanted microtissues. Ceria nanoparticle treatment of implanted microtissues reduced lipid peroxidation and protected neuronal density around the implant, while restoring functional connectivity and community structure of microcircuits. These results suggest that the innate immune system, via ROS-mediated neuroinflammation, may play a critical role in the foreign-body response to neural implants, independent of the systemic immune response and initial implantation injury. Further, we demonstrate for the first time the potential for antioxidant ceria nanoparticles to treat neuroinflammation at the device-tissue interface.

## Results

### Development of a 3D in vitro model of the device-tissue interface

We developed a 3D in vitro model of the device-tissue interface to test the effect of ceria nanoparticles without the need for an advanced delivery method to bypass the blood brain barrier. We employed the use of a primary 3D in vitro model of the cortex ^50^. Briefly, cortical tissue was collected from postnatal day 1 rats (**Figure 1A**) and dissociated into a single cell suspension (**Figure 1B**). To visualize a larger area of the wire implant surface, cells were seeded along with wire into an agarose microwell (**Figure 1C**). At DIV1, cells produce an endogenous extracellular matrix, forming a microtissue (**Figure 1D**). The placement of the agarose microwell close to the bottom of the plate placed the microtissue within the working distance of a confocal 30x objective, allowing for live imaging of oxidative stress, functional calcium activity, and structural neuronal networks in microtissues. Oxidative stress data was collected at DIV14 by measuring oxidation over a 30-minute incubation period (**Figure 1E**) using a fluorescent lipid peroxidation reporter (Image IT Lipid Peroxidation Kit, **Figure 1F**). Functional and structural data was obtained through the application of a genetically encoded calcium indicator (GECI) with an mRuby control tag. AAV mediated delivery of fluorescent proteins was performed at DIV 1 (**Figure 1G**), producing stable fluorescence by DIV 14. 3D neuronal network structure (red, **Figure 1H**) Calcium activity (green, **Figure 1I**) and were imaged at DIV 14, 16 and 18.

**Figure 1.**
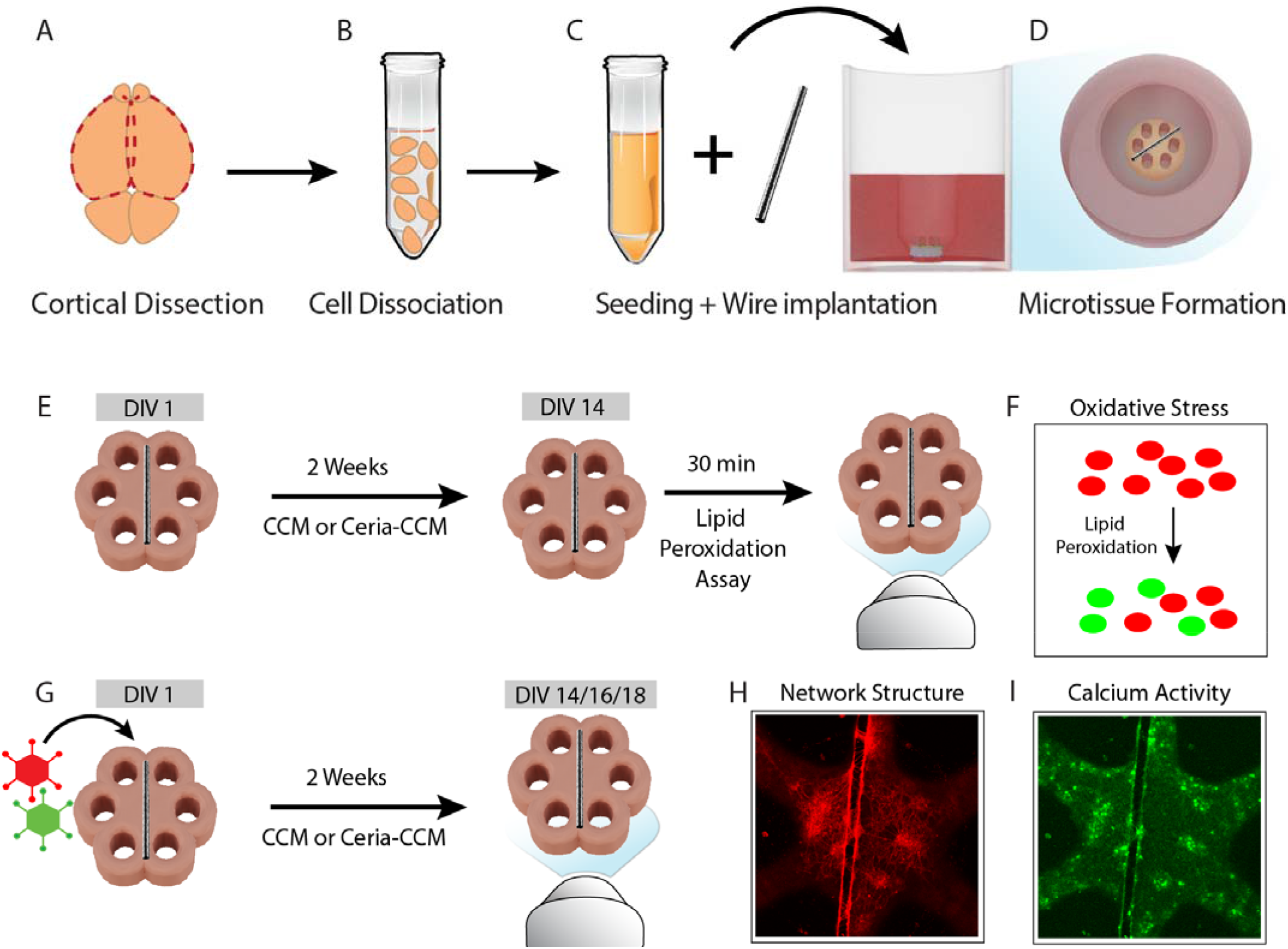
Development of an in vitro device-tissue interface for measuring oxidative stress, functional activity, and structural changes around implanted wires. (A) Cortices were dissected away from P1 rat brains. (B) Primary tissue was dissociated into a single cell suspension by papain. (C) Cells were seeded alongside a tungsten wire into an agarose microwell. (D) Cells formed a microtissue with an embedded microwire. (E) Oxidative stress assays were performed after 14 days of culturing. (F) Image-IT kit shifts live cell fluorescence from red to green in the presence of lipid peroxidation over 30 minutes. (G) AAV mediated fluorescent protein delivery was performed at day one in vitro, and transfected microtissues were imaged on DIV 14, 16 and 18. (H) Network structure was imaged in the red channel with a 30x oil object for high resolution volumetric data. (I) Calcium activity was recorded in the green channel with a 10x objective for 4min with a 15Hz sampling rate.

### Local lipid peroxidation occurs at the device-tissue interface

Oxidative stress around implants is an important hallmark of the in vivo chronic foreign body response^17^, and thus, an important feature to evaluate in vitro. To characterize oxidative stress around the implant in vitro, we used a fluorescent reporter of lipid peroxidation (Image-iT Lipid Peroxidation Kit) in which a fluorescent reagent shifts from red to green when the dye undergoes oxidation. A lower red/green fluorescence ratio indicates a higher degree of lipid peroxidation. Lipid peroxidation of control microtissues was assessed by measuring the red/green fluorescence ratio over the entire center region of the tissue by using the “whole-tissue” region of interest (ROI) (gray circle, **Figure 2A**). Lipid peroxidation of wire implanted microtissues was assessed by measuring the red/green ratio as a function of distance from the implant surface by using 50µm binned ROIs placed from 0-400µm from the implant surface (pink, **Figure 2B**). The red/green ratio within 200µm of the surface of the wire was significantly lower than the control, indicating increased lipid peroxidation at the device-tissue interface (50µm, p=0.408; 100µm, p=0.0207; 150µm, p=0.0198; 200µm, p=0.0306; 250µm p=0.0825; 300µm, p=0.1973; 350µm, p=0.4378; 400µm, p=0.6462; **Figure 2C**). These results suggest that oxidative stress occurs within 200µm of the device-tissue interface in vitro, without an implantation wound or chronic vascular instability associated with oxidative stress in the in vivo environment.

**Figure 2.**
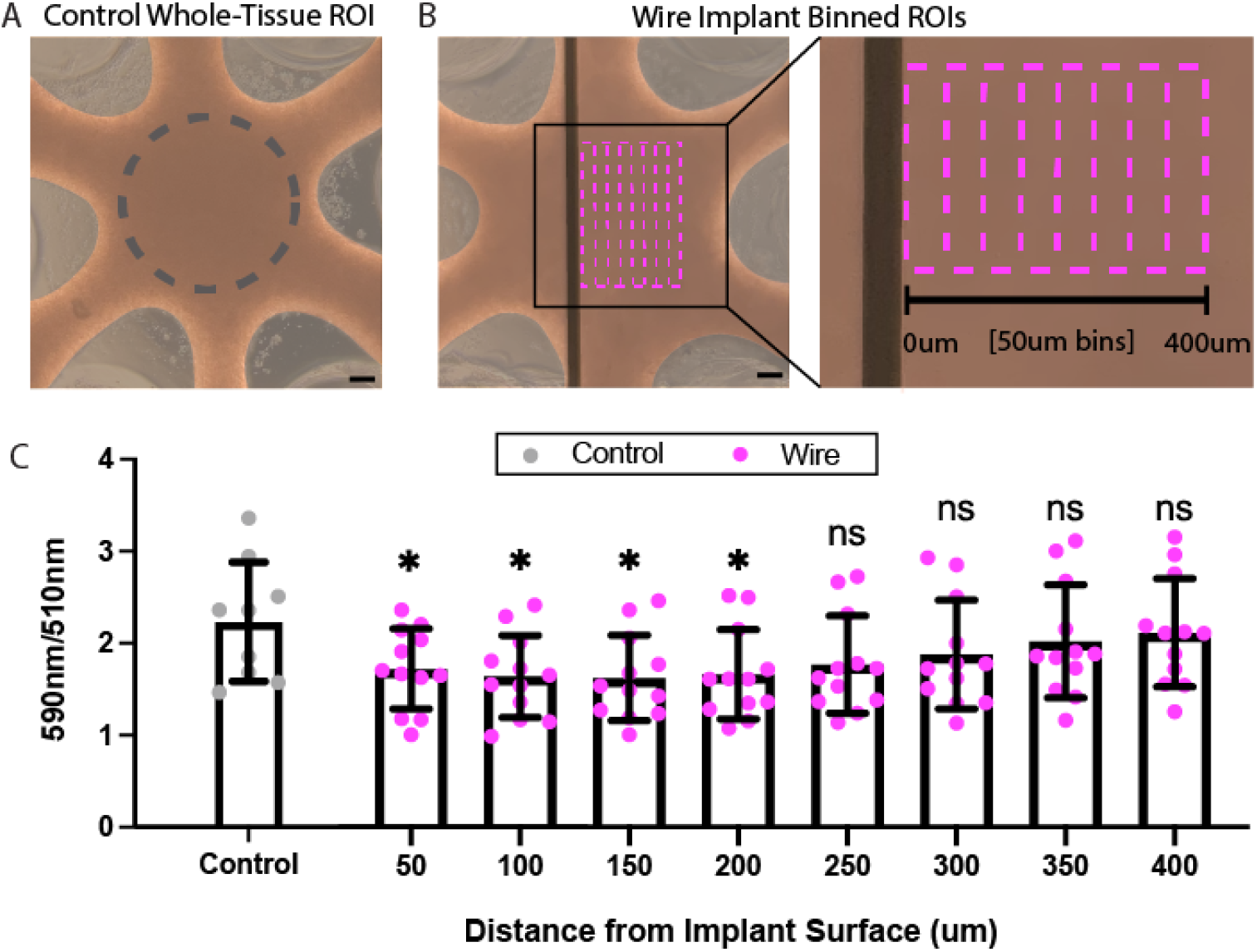
Lipid peroxidation occurs locally around implants. (A) Lipid peroxidation of control microtissues was determined by measuring the red/green ratio over the entire center of the microtissue (gray dashed circle). (B) Lipid peroxidation of wire implanted microtissues was determined by measuring the red/green ratio as a function of distance from the surface of the wire (0-400 µm from the wire in 50 µm bins). (C) The red/green ratio within 200 microns of the wire was significantly lower than the whole-tissue control, indicating a significant increase in lipid peroxidation around the implant compared to unimplanted tissue (50µm, p=0.408; 100µm, p=0.0207; 150µm, p=0.0198; 200µm, p=0.0306; 250µm p=0.0825; 300µm, p=0.1973; 350µm, p=0.4378; 400µm, p=0.6462). Scale bars = 100µm. *p<0.05.

### Implants reduce neuronal density at the device-tissue interface

We further investigated changes at the device-tissue interface and the effects of ceria treatment by examining the live network structure. Live neuronal morphology was achieved by AAV-mediated transgene delivery of an mRuby fluorescent tag (AAV1-hSyn1-mRuby2-GSG-P2A-GCaMP6s-WPRE-pA, AddGene), which was expressed in neurons and imaged at DIV 14, 16, and 18 (**Figure 3A**). Three-dimensional 30x fluorescent images of the neuronal structure in the center of the microtissue were binarized using Otsu’s thresholding method (**Figure 3B**) and neuronal volume fraction was calculated by the sum of positive voxels/total voxels within the ROI (control ROI example image, **Figure 3C**).

**Figure 3.**
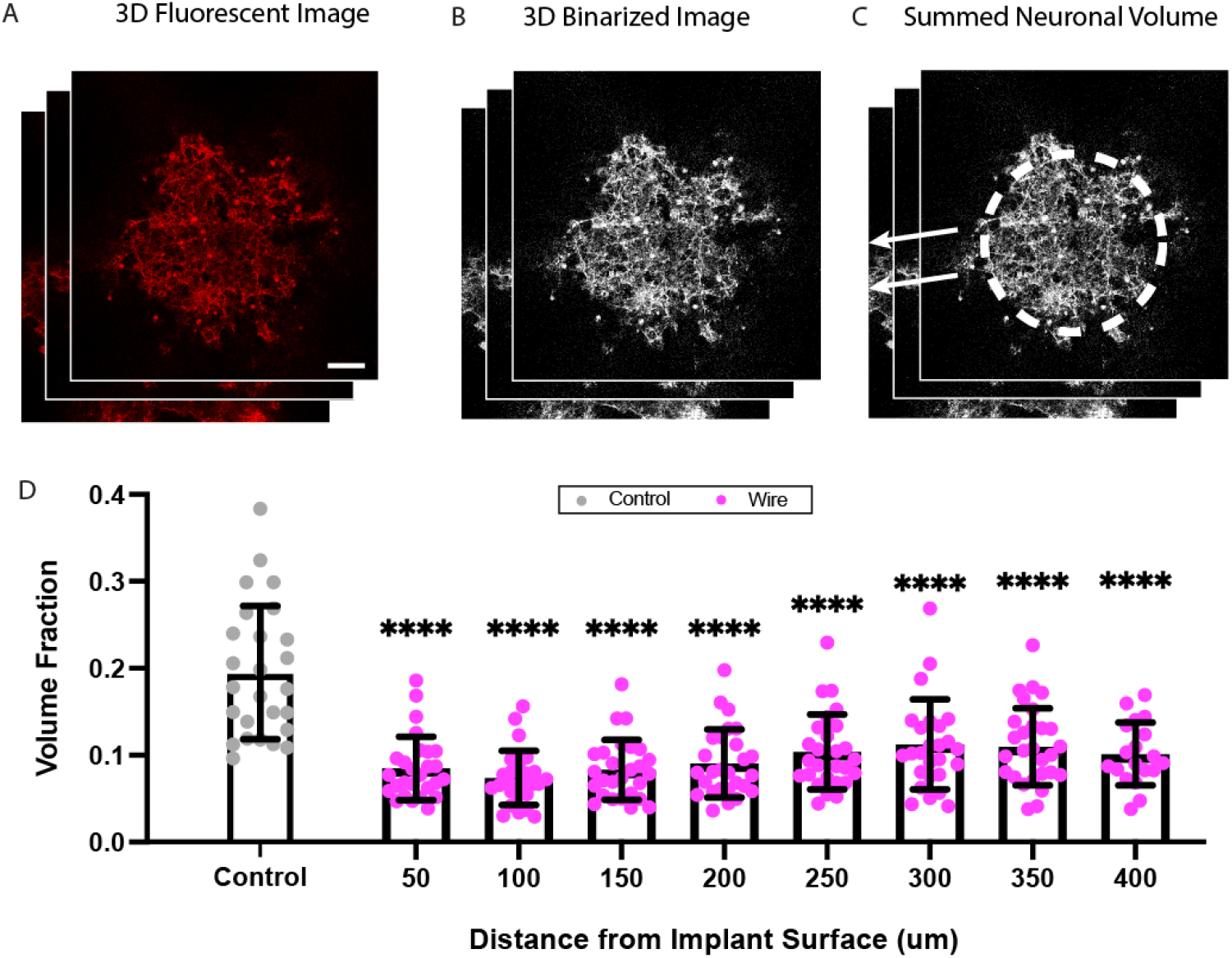
Neuronal density is reduced around implants. (A) 3D 30x volume images of the structural mRuby tag in the center of the microtissue were collected from live from DIV 14, 16 and 18. (B) Fluorescent images were binarized using Otsu’s thresholding method in FIJI. (C) Neuronal density was determined by measuring the volume fraction of white voxel from binarized volumes divided by the total number of voxels within the ROI (control example ROI gray dashed circle). (D) Neuronal volume fractions 400µm from the surface of the implanted wires (pink) were significantly lower than controls (gray) (50µm, p<0.0001; 100µm, p<0.0001; 150µm, p<0.0001; 200µm, p<0.0001; 250µm p<0.0001; 300µm, p<0.0001; 350µm, p<0.0001; 400µm, p<0.0001). Scale bars = 100µm. ****p<0.0001.

We found that local neuronal density, as measured by the neuronal volume fraction within each 50µm bin from 0-400µm from the edge of the wire implant, were significantly reduced from the neuronal density in control microtissues (50µm, p<0.0001; 100µm, p<0.0001; 150µm, p<0.0001; 200µm, p<0.0001; 250µm p<0.0001; 300m, p<0.0001; 350µm, p<0.0001; 400µm, p<0.0001; **Figure 3D**). Overall, these data indicate that wire implants significantly disrupt neuronal network structure.

### Implants increase functional connectivity and disrupt community structure of in vitro microcircuits

In order to assess the functional effects of wire implantations, we used AAV-mediated transgene delivery of a GCaMP6s calcium reporter (AAV1-hSyn1-mRuby2-GSG-P2A-GCaMP6s-WPRE-pA, AddGene) and recorded calcium activity in the tissue surrounding implants from DIV14-18. We used a mathematical method called graph theoretical analysis (GTA) to describe the complex functional interactions within a network^51,52^. Application of GTA to a graph made up of single cell “nodes” is a characterization of network behavior of “microcircuits”^53,54^, which are sensitive to subtle changes in network dynamics that precede gross histological and behavioral deficits in other neuroinflammatory and neurodegenerative disease states^55–58^. Examination of microcircuit dynamics is particularly well suited to the device-tissue interface, as the neuroinflammatory response is localized to the device and likely does not extend to multiple brain regions at the “mesocircuit” level^59^. We applied GTA to time-series fluorescent calcium traces to characterize three metrics of node-pair interactions: correlation, clustering, and path length. Correlation, which represents the how similar calcium activity between two nodes are overtime, was calculated by averaging the Pearson cross correlation coefficient of every node-pair in the microtissue. Clustering is a metric of “small world” network organization, which measures functional connectivity between triplets of nodes^31,51,60^. Path length, an indicator of the efficiency of transmission between nodes, is inversely proportional to correlation, Both path length and clustering were calculated using the network analysis toolkit from Dingle et al^31^.

We then assessed whole-tissue functional connectivity analysis, which included all cells in the microcircuit regardless of position in relation to the wire implantation. Wire implanted microtissues exhibited significantly higher whole-tissue correlations compared to controls (Wire, 0.6703 ± 0.1018; Control, 0.5948 ± 0.08139; p=0.0040; **Figure 4A**). Wire implanted microtissues also showed a significant increase in clustering (Wire, 0.6617 ± 0.1053; Control, 0.5885 ± 0.08258; p =0.0064; **Figure 4B**), but not path length (Wire,1.760 ± 0.3435; Control, 1.921 ± 0.3631; p =0.1005; **Figure 4C**). These data suggest substantial functional reorganization of neural networks surrounding implants.

**Figure 4.**
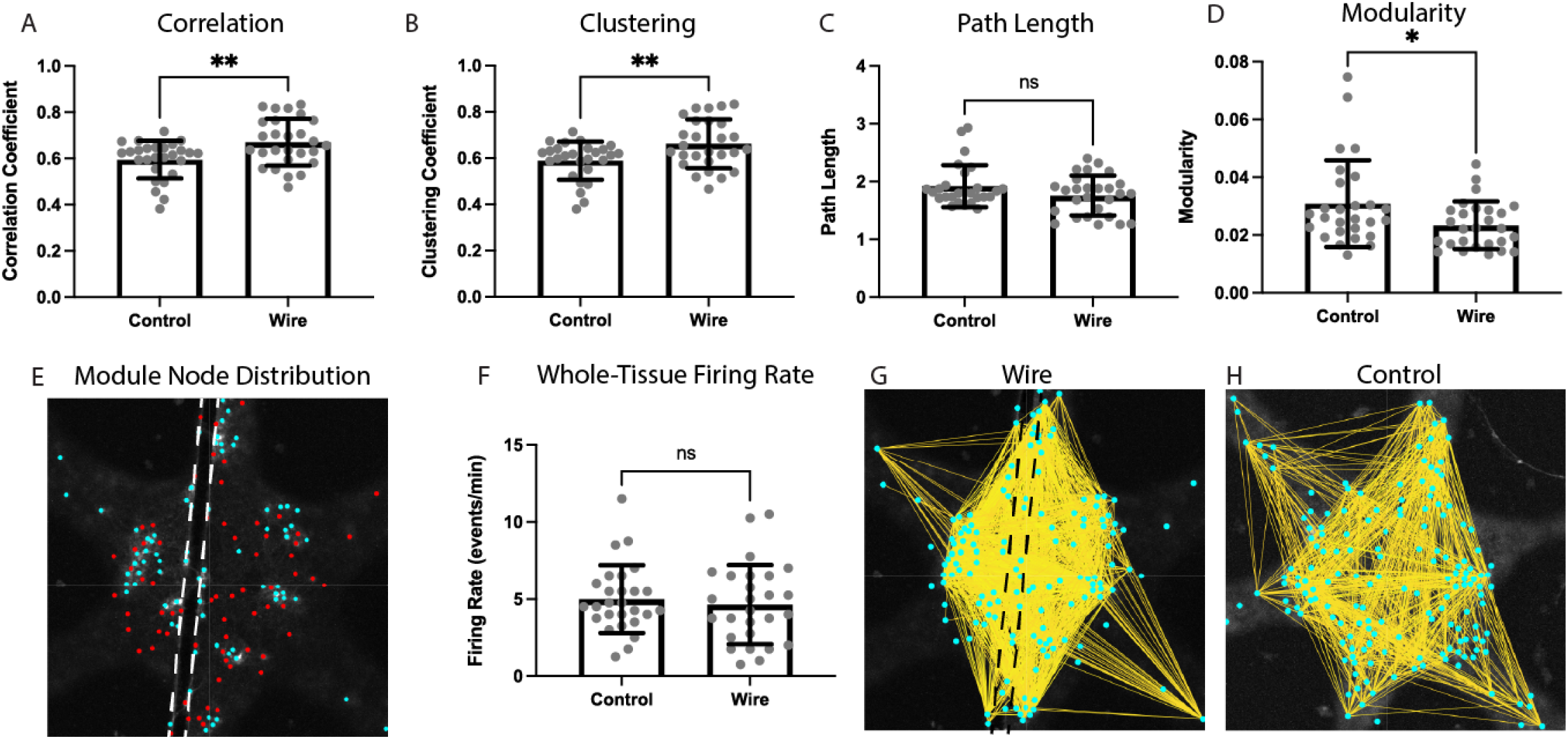
Wire implants increase whole tissue functional connectivity and disrupt community structure. (A) Correlation of wire implanted microtissues are significantly higher than controls (p = 0.0040). (B) Clustering significantly increased in wire implanted samples compared to controls (p = 0.0064). (C) Path length showed no significant difference between controls and wire implanted microtissues (p = 0.1005). (D) Modularity was significantly decreased in wire implanted samples (p = 0.0272). (E) A representative microtissue shows that spatial distribution of modules was not related to wire position indicated by dashed white lines. (F) Whole tissue firing rate was not significantly affected by wire implantation (p =0.5905). Representative correlational connectomes visually show the significant increase in high correlation values (>0.9) in wire implanted microtissues (G, wire position indicated by black dashed lines) compared to unimplanted controls (H). *p<0.05, **p<0.01.

Finally, we assessed modularity, a metric of network segregation associated with learning and memory processes. We calculated modularity using a hierarchical clustering method which identifies groups of highly correlated cells^31,61^. Wire implantation significantly reduced modularity of the network compared to controls (Wire,0.02340 ± 0.008260; Control, 0.03088 ± 0.01499; p =0.0272; **Figure 4D**). Interestingly, the spatial distribution of modules was not related to implant position (**Figure 4E**). These results indicate that implantation disrupts critical community structure of microcircuits beyond the local network directly around the implant.

The increase in functional connectivity had no significant effect on whole tissue firing rate (Wire, 4.639 ± 2.564; Control, 4.991 ± 2.198; p =0.5905; **Figure 4F**). Such changes in functional connectivity can be visualized with a correlational connectome, which shows a clear increase in highly correlated connections (>0.9) in wire implanted (**Figure 4G**) compared to control (**Figure 4H**) microtissues.

### Optimization of ceria nanoparticle coatings for treatment of oxidative stress

We then sought to investigate the efficacy of antioxidant ceria nanoparticles in protecting the device-tissue interface and, in turn, examine the role of ROS in mediating innate neuroinflammation. Ceria nanoparticles are cerium(III) rich at the particle interface, which subsequently results in more oxygen vacancies in smaller nanocrystals^42,43^. This makes ceria nanoparticles excellent oxygen buffers, which can easily absorb and release oxygen during high temperature catalytic processes^62^. They are also potent antioxidants in water, capable of neutralizing large amounts of many types of reactive oxygen species (ROS). During this process cerium(III) is oxidized to cerium(IV), but over days and weeks the fully oxidized particles eventually return to their cerium(III)-rich state^44^. Such perpetual, temperature-stable antioxidants are ideal for treating chronic neuroinflammation at the device-tissue interface. However, application of nanoscale ceria to biological tissues requires a polymer coating to ensure stability and prevent nanoparticle aggregation.

Here, we synthesized catecholamine-functionalized polyethylene glycol (PEG) and used it to coat uniform and crystalline ceria nanoparticles (d = 5.1 nm ± 1.0 nm) prepared in organic media (**Figure 5A**). We tested three catecholamine-functionalized PEG coatings with molecular weights of 5kDa (**Figure 5B**), 10kDa (**Figure 5C**), and 40kDa (**Figure 5D**). The catecholamine bonds to the surface cerium atoms resulting in biocompatible ceria nanoparticles in water, which can be visualized by cryo-TEM (**Figure 5E**). Increases in the PEG molecular weight resulted in increased hydrodynamic diameter in both water and PBS (**Figure 5F**), as well as decreased graphing density (**Figure 5G**). Additionally, we found that the attachment of the catecholamine at the surface of the nanoparticle increased the antioxidant capacity, making the high graphing density 5k PEG-nitro-DOPA coated sample the most effective antioxidant. MTT toxicity testing of 5k PEG-nitro-DOPA coated ceria showed no acute toxicity to primary neurons at 14ppm (Supplemental **Figure S1**). As such, we chose to use the 5k PEG-nitro-DOPA coated ceria for all subsequent biological efficacy testing.

**Figure 5.**
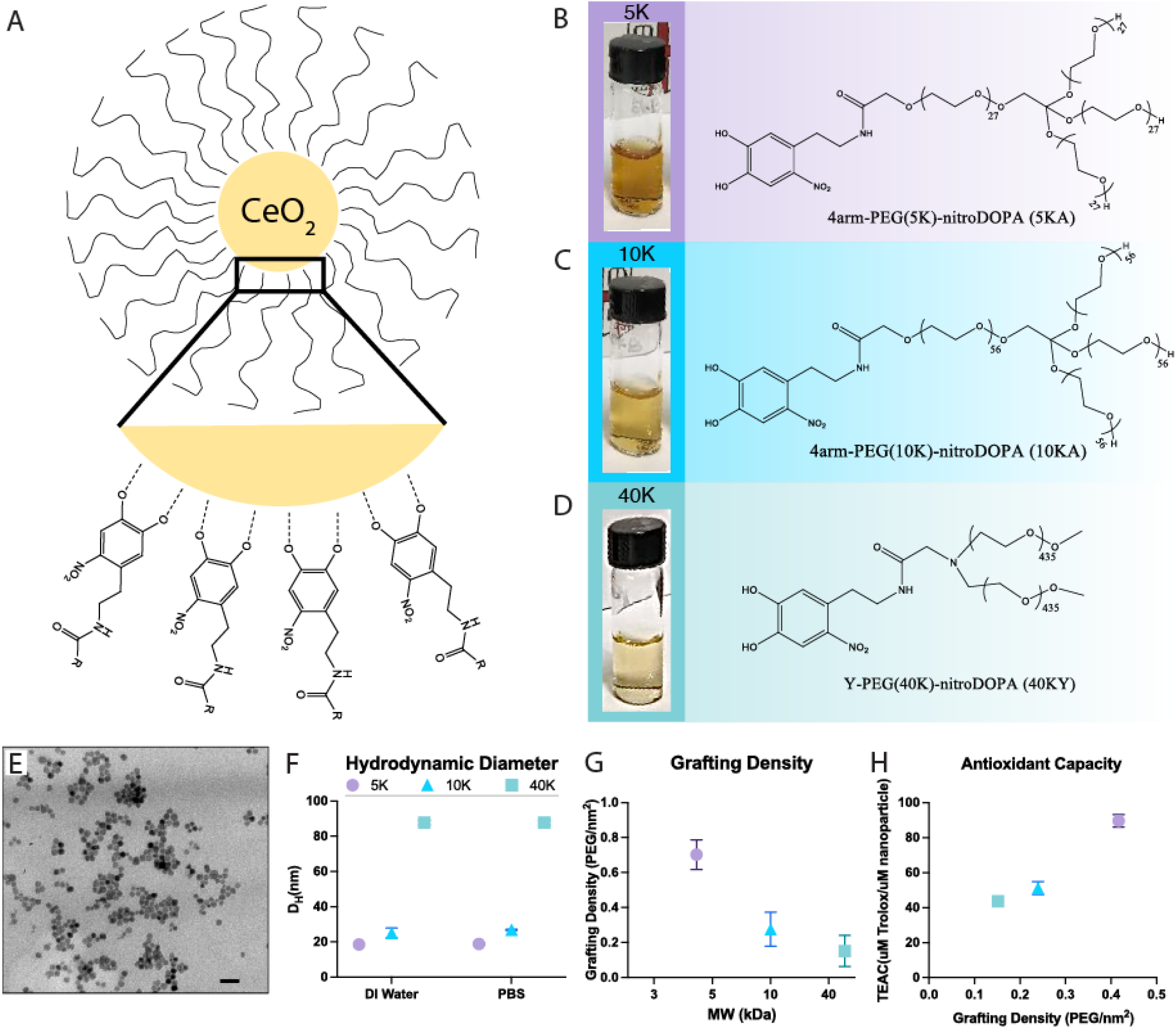
The 5K-PEG-nitro-DOPA coating has optimal characteristics for biological applications. (A). Scheme of ceria nanoparticles coated by nitroDOPA-PEG polymers. (B) Photo of nitroDOPA-PEG(5K) functionalized ceria nanoparticles in water ([Ce]=500 ppm). (C) Photo of nitroDOPA-PEG(10K) functionalized ceria nanoparticles in water ([Ce]=500 ppm). (D) Photo of nitroDOPA-PEG(40K) functionalized ceria nanoparticles in water ([Ce]=500 ppm). (E) Cryo-TEM photo of the nitroDOPA-PEG(10K) coated ceria nanoparticles, scare bar = 20nm. (F) Mean and relative variance of the size distributions of the ceria nanoparticles measured by dynamic light scattering (DLS) in water and PBS buffer. (G) Grafting densities of nitroDOPA-PEG coated nanoparticles decrease while PEG chain lengths increase. Thermogravimetric analysis (TGA) was collected on the Mettler Toledo TG50 Thermogravimetric with analyzer using an alumina crucible. All samples started with a mass at 3-5 mg. Samples were first heated to 90 °C and held for half an hour to get rid of humidity. The heating rate of the analysis was 20 °C/min between 100 and 950 °C, under air atmosphere with a flow rate of 80 ml/min. All samples were tested three times. (H) Evaluation of antioxidant capacity for ceria nanoparticles by Trolox (6-hydroxy-2,5,7,8-tetramethylchroman-2-carboxylic acid) equivalent antioxidant capacity (TEAC) assay using sodium fluorescein (FL) as the fluorescent probe. TEAC values of ceria nanoparticles with different PEG coatings were compared to their grafting densities.

### Ceria treatment mitigates oxidative stress at the device-tissue interface

After optimization of the ceria formulation, we then characterized the effects of ceria nanoparticles on neuroinflammation around implanted wires. This investigation will provide insight into the role of ROS in the in vitro neuroinflammatory response, as well as an indication of biological efficacy of ceria in protecting the device-tissue interface. Cells were exposed to a 14ppm ceria solution diluted in media every other day for 14 days in vitro before performing the lipid peroxidation assay.

We found that ceria treatment mitigated oxidative stress in wire implanted microtissues, with a significant increase in the red/green fluorescence ratio in ceria treated wire implanted samples (purple) from 0-400µm compared to untreated wire implants (pink) (50µm, p=0.0091; 100µm, p=0.0107; 150µm, p=0.0067; 200µm, p=0.0117; 250µm p=0.0538; 300µm, p=0.1306; 350µm, p=0.3251; 400µm, p=0.5679; **Figure 6**, significance indicated above brackets). While wire implantation significantly increased oxidative stress compared to unimplanted controls with 200µm of the wire (50µm, p=0.408; 100µm, p=0.0207; 150µm, p=0.0198; 200µm, p=0.0306; 250µm p=0.0825; 300µm, p=0.1973; 350µm, p=0.4378; 400µm, p=0.6462; **Figure 6**, significance indicated above pink data points), ceria treatment returned oxidative stress levels back to that of unimplanted controls within 400µm of the wire surface (50µm, p=0.5729; 100µm, p=0.3666; 150µm, p=0.4019; 200µm, p=0.4808; 250µm p=0.5097; 300µm, p=0.7006; 350µm, p=0.8685; 400µm, p=0.9426; **Figure 6**, significance indicated above purple data points). These data indicate that ceria treatment mitigates oxidative stress at the device-tissue interface.

**Figure 6.**
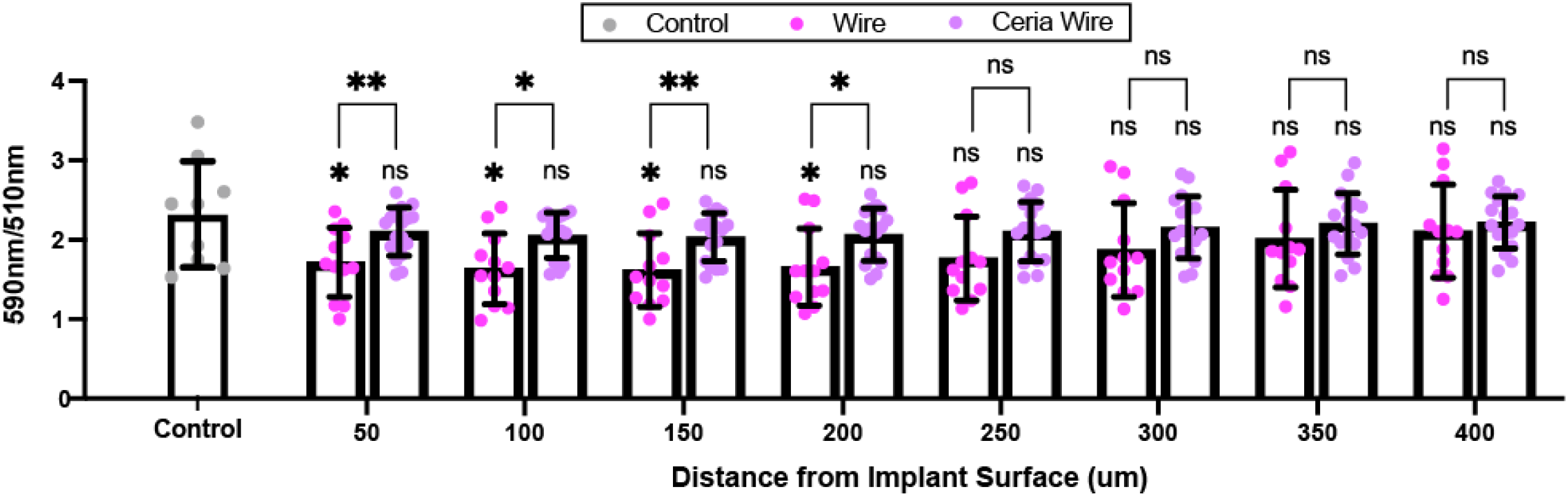
Ceria nanoparticles significantly reduce oxidative stress around implants. Ceria treated wire implants (purple) exhibit significantly increased red/green fluorescence over untreated wire implants (pink) (50µm, p=0.0091; 100µm, p=0.0107; 150µm, p=0.0067; 200µm, p=0.0117; 250µm p=0.0538; 300µm, p=0.1306; 350µm, p=0.3251; 400µm, p=0.5679; significance indicated above brackets). Wire implants (pink) exhibit significantly decreased red/green fluorescence from 0-200µm compared to unimplanted controls (gray) (50µm, p=0.408; 100µm, p=0.0207; 150µm, p=0.0198; 200µm, p=0.0306; 250µm p=0.0825; 300µm, p=0.1973; 350µm, p=0.4378; 400µm, p=0.6462; significance indicated above pink data points). Ceria treated wire implants (purple) show no significant difference compared to untreated unimplanted controls (gray) (50µm, p=0.5729; 100µm, p=0.3666; 150µm, p=0.4019; 200µm, p=0.4808; 250µm p=0.5097; 300µm, p=0.7006; 350µm, p=0.8685; 400µm, p=0.9426; significance indicated above purple data points). *p<0.05, **p<0.01.

### Ceria treatment reduces structural changes around implants

As oxidative damage can directly affect structural neural networks, we assessed the effect of ceria treatment on neuronal density. Ceria treatment of wire implanted samples (purple) showed significantly higher neuronal density around electrodes compared to untreated wire implants (pink) within 400µm of the implant surface (50µm, p=0.0002; 100µm, p<0.0001; 150µm, p<0.0001; 200µm, p<0.0001; 250µm p=0.0001; 300µm, p=0.0003; 350µm, p<0.0001; 400µm, p<0.0001; **Figure 7A**, significance indicated above brackets). While wire implants exhibited significant density loss within 400µm of the implant compared to unimplanted controls (50µm, p<0.0001; 100µm, p<0.0001; 150µm, p<0.0001; 200µm, p<0.0001; 250µm p<0.0001; 300µm, p<0.0001; 350µm, p<0.0001; 400µm, p<0.0001; **Figure 7A**, significance indicated above pink data points), neuronal density loss in ceria treated samples was reduced within 250µm of the implant and fully mitigated from 250-400µm (50µm, p=0.0007; 100µm, p=0.0044; 150µm, p=0.0124; 200µm, p=0.0141; 250µm p=0.0126; 300µm, p=0.2662; 350µm, p=0.5218; 400µm, p=0.1270; **Figure 7A**, significance indicated above purple data points). Representative images of an unimplanted control (**Figure 7B**), wire implanted sample (**Figure 7C**), and ceria treated wire implanted sample (**Figure 7D**) show a clear reduction in neuronal density around untreated wire implants and protection of neuronal density around ceria treated implants. Overall, these data indicate that ceria treatment prevents structural damage to neuronal networks caused by neuroinflammation around implants.

**Figure 7.**
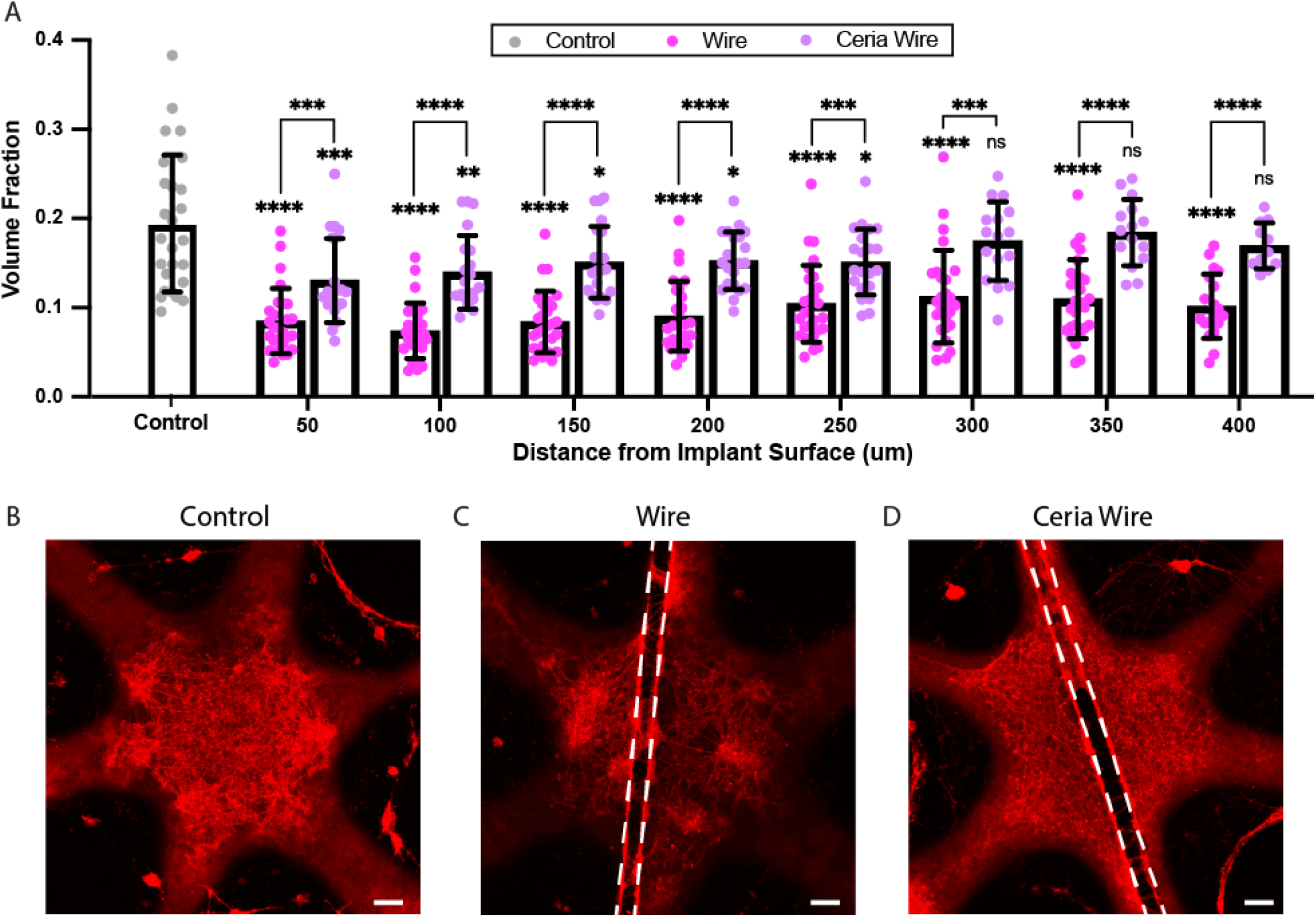
Neuronal density loss around implants is mitigated by ceria treatment. (A) Neuronal volume fractions of wire implants (pink) were significantly lower than unimplanted controls (gray) (50µm, p<0.0001; 100µm, p<0.0001; 150µm, p<0.0001; 200µm, p<0.0001; 250µm p<0.0001; 300µm, p<0.0001; 350µm, p<0.0001; 400µm, p<0.0001, significance indicated above pink data points). Ceria treated wire implants (purple) exhibited significant increases in neuronal volume factions over untreated wire implants (50µm, p=0.0002; 100µm, p<0.0001; 150µm, p<0.0001; 200µm, p<0.0001; 250µm p=0.0001; 300µm, p=0.0003; 350µm, p<0.0001; 400µm, p<0.0001; significance indicated above brackets), reducing neuronal density loss from 0-250µm and restoring neuronal densities to that of unimplanted controls from 250-400µm (50µm, p=0.0007; 100µm, p=0.0044; 150µm, p=0.0124; 200µm, p=0.0141; 250µm p=0.0126; 300µm, p=0.2662; 350µm, p=0.5218; 400µm, p=0.1270; significance indicated above purple data points). The loss of neuronal density around wire implants and subsequent protection of neuronal density after ceria treatment can be seen visually in representative images of (B) unimplanted controls, (C) wire implants, and (D) ceriatreated wire-implants. Scale bars = 100µm. *p<0.05, **p<0.01, ***p<0.001, ****p<0.0001.

### Ceria treatment restores functional connectivity and protects community structure in implanted microtissues

We characterized the effects of antioxidant ceria nanoparticles in protecting neurons against the functional connectivity changes caused by wire implantation. Such characterization of functional network activity around implants in after removal of excess ROS will indicate if disruption of functional connectivity is resultant from ROS-mediate neuroinflammation. To achieve this, microtissues were treated with ceria nanoparticles diluted to 14ppm in media every other day for the 18 days in vitro, replacing 500uL of liquid in each media change. Ceria treatment significantly decreased correlation in wire implanted microtissues (Ceria Wire,0. 5420 ± 0.1380; Wire, 0.6703 ± 0.1018; p =0.0002; **Figure 8A**).

**Figure 8.**
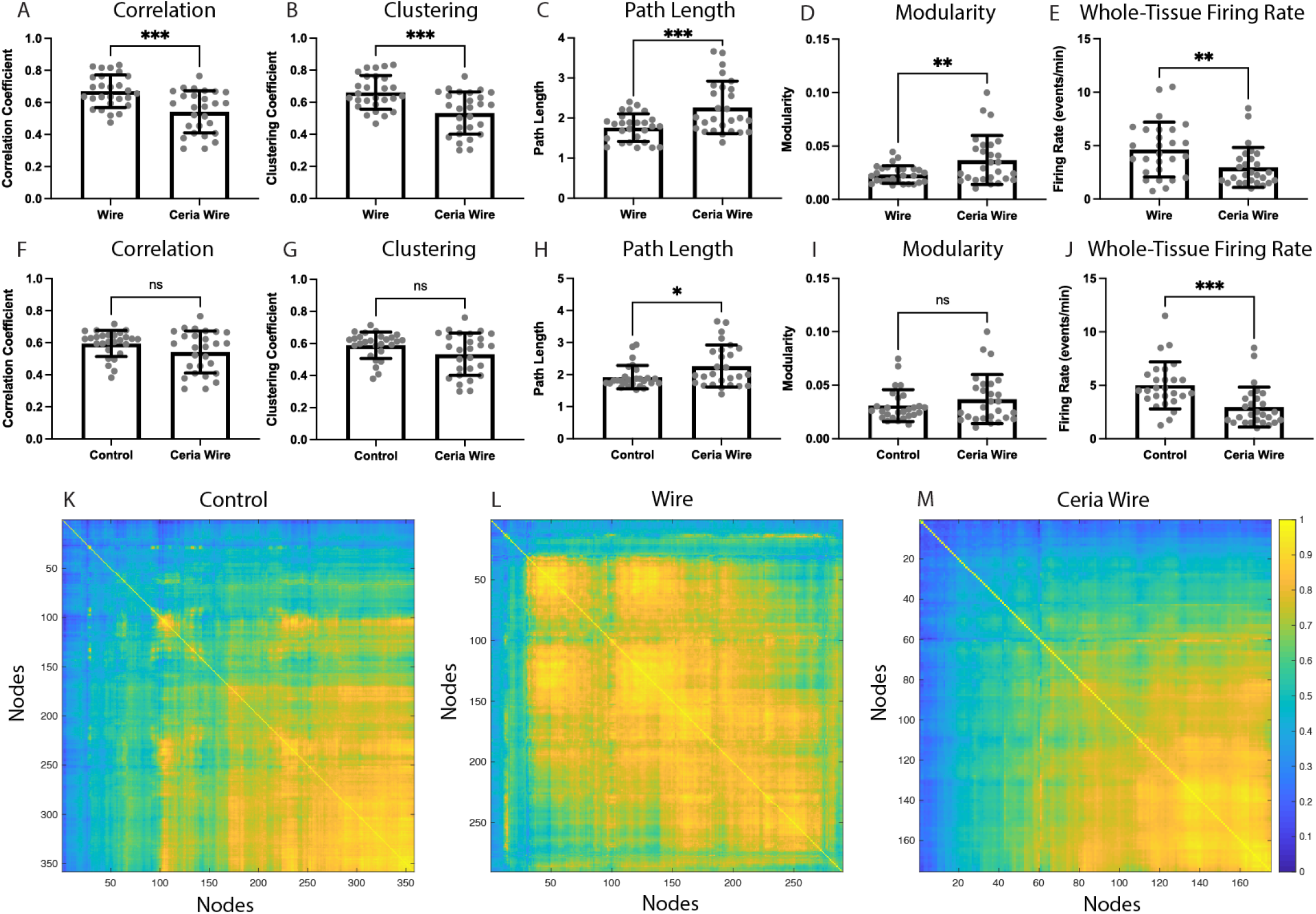
Ceria treatment restores functional connectivity and protects community structure around implants. Ceria treatment significantly reduced correlation (A, p= 0.0002) and clustering (B, p = 0.0002), while reducing path length (C, p = 0.0008) around wire implants. Ceria treatment strengthened community structure through increased modularity (D, p = 0.0056). Whole tissue firing rate was significantly decreased ceria wire implants compared to untreated wire implants (E, p = 0.0083). Correlation of ceria treated wire implanted microtissues is not significantly different from controls (F, p = 0.0819). Clustering also showed no significant difference between ceria treated wires and controls (G, p = 0.0692). Path length showed a small, but significant increase in ceria treated wire samples from unimplanted controls (H, p = 0.0441). Ceria treatment restored modularity of the network in implanted tissues to control levels (I, p = 0.2545). Whole tissue firing rate was significantly decreased by ceria treatment in wire implanted samples compared to unimplanted control microtissues (J, p = <0.0001). Correlograms from representative (K) control, (L) wire implanted, and (M) ceria treated wire implanted microtissues. *p<0.05, **p<0.01, ***p<0.001.

Ceria treated microtissues similarly exhibited reduced clustering around wire implants (Ceria Wire,0.5328 ± 0.1317; Wire, 0.6617 ± 0.1053; p =0.0002; **Figure 8B**), and increased path length (Ceria Wire, 2.266 ± 0.6559; Wire, 1.760 ± 0.3435; p =0.0008; **Figure 8C**). These results suggest that ceria nanoparticles reduce the functional connectivity changes at the device-tissue interface. Additionally, ceria treatment improved the strength of community structure around implants, with a significant increase in modularity (Ceria Wire, 0.03697 ± 0.02293; Wire, 0.2340 ± 0.008260; p =0.0056; **Figure 8D**). Remarkably, ceria treatment also significantly reduced whole tissue firing rate (Ceria Wire, 2.963 ± 1.872; Wire, 4.639 ± 2.564; p =0.0083; **Figure 8E**).

We then compared functional connectivity around ceria treated wires to that of unimplanted controls. Correlations of ceria treated wire implanted microtissues were not significantly different from correlations of unimplanted controls (p = 0.0819, **Figure 8F**). Additionally, clustering of ceria treated wire implanted microtissues was not significantly different from unimplanted controls (p = 0.0692, **Figure 8G**). Interestingly, path length of ceria treated wire implants showed a small statistical difference from controls (p = 0.0441, **Figure 8H**). These data, while on the edge of significance, suggest that ceria treatment may restore functional connectivity of wire implanted microtissues closer to that of unimplanted controls.

Further investigation of the network revealed that ceria treatment restored modularity of wire implanted microtissue to control values (p = 0.2545, **Figure 8I**), indicating that ceria treatment protected community structure around implants. Additionally, treatment significantly reduced the whole-tissue firing rate compared to controls (p = 0.0001, **Figure 8J**), suggesting that ceria treatment of the whole tissue via media delivery may interfere with synchronous network activities. Correlograms from representative microtissues display the clear increase in correlation from wire implantation and subsequent mitigation of the functional changes with ceria treatment (**Figure 8K-M**). Collectively, these results indicate that ceria nanoparticles protect neural function around the device-tissue interface in this in vitro model.

## Discussion

We developed a 3D in vitro model of the device-tissue interface to better understand the independent contributions of the innate immune system on the neuroinflammatory response to devices. We found that an isolated innate immune response, in the absence of vascular derived immune drivers, was capable of recapitulating important hallmarks of the in vivo response, including increased oxidative stress and decreased neuronal density within 250µm of the implant surface. These symptoms were accompanied by significant changes in functional connectivity and community structure. To understand the role of ROS-mediated oxidative damage in this innate neuroinflammatory response, we tested the effects of antioxidant ceria nanoparticles, optimized for hydrodynamic diameter and antioxidant capacity, on the symptoms of neuroinflammation. We found that ceria nanoparticles reduced oxidative stress, increased neuronal density, and restored metrics of functional connectivity and community structure.

Increased oxidative stress and decreased neuronal density in an avascular model of the vitro device-tissue interface indicate the presence of a distinct innate immune response to microwire implants. These results suggest that, while vascular instability is a significant contributor to the chronic inflammatory response in vivo, it is not necessarily the only driver of chronic inflammation around implants. Our work is the first to show oxidative stress and neuronal density loss directly around implants in an in vitro environment, improving physiological relevance over previous 2D models^28^. Additionally, we found that tis tissue reaction occurs in the absence of an acute stab wound from implantation, suggesting that innate glia may be sensing and responding to chemical or mechanical cues from the implant, as demonstrated elsewhere ^63–66^. Increases in functional connectivity and disruption of community structure in the wire-implanted microtissues were consistent with LPS-induced neuroinflammatory responses both in vitro and in vivo, alluding to the fact that functional and structural changes to neurons and their synaptic behaviors may be mediated by glial dysregulation^50,55^. Remarkably, we found that while symptoms of oxidative tissue damage occurred locally at the implant surface, changes in functional network dynamics affected the whole microtissue, revealing that localized neuroinflammation impacts the surrounding microcircuitry. The changes in neural network dynamics reveal a significant “observer effect” from exogenous material implantation and has broad implications for the use of invasively recorded data in studying normal neural activity. Further, the influence of the multiphasic neuroinflammatory state on functional connectivity may contribute, in part, to the long-term signal drift of recordings that results in inconsistent decoding for prosthetic control^67,68^.

We demonstrate the use of ceria nanoparticles for reducing oxidative stress, increasing neuronal density, and reducing functional connectivity in wire implanted samples. The restoration of functional connectivity after antioxidant treatment suggests that the structural and functional changes around electrodes are likely downstream symptoms of oxidative stress. In vivo antioxidant treatments have shown similar reductions in oxidative stress and increases in neuronal density. Additionally, these results indicate that antioxidant ceria nanoparticles may be an efficacious drug treatment for protecting neural function at the device-tissue interface. While these acute treatment results are promising, the true advantage of employing a recyclable antioxidant like ceria nanoparticles lies in the long-treatment capacity, which will require further testing.

While ceria treatment was neuroprotective under oxidative stress conditions, ceria treatment of unimplanted tissues showed significant changes in neuronal density and functional connectivity compared to untreated controls (**Supplemental Figures S2, S3, S4, S5**). As ROS is an important signaling molecule involved in normal synaptic regulation^69,70^, neutralization of ROS in healthy tissues with a normal oxidative load can interfere with functional connectivity. We found that ceria affected the whole tissue firing rate of microtissues, in both wire-implanted and non-implanted microtissues, which may be an example of antioxidant overdosing interfering with synchronized bursts (**Supplemental Figure S4**). We hypothesize that the ideal antioxidant treatment strategy should utilize an adaptive local delivery method to treat only the local device-tissue interface when the oxidative load becomes too large. Future work will also explore other metrics of neuroimmune interactions, through live cell morphology imaging to assess real-time microglial activity, to a variety of implanted materials, including glass and parylene coated electrodes.

While the 3D in vitro microenvironments presented here closely resemble their in vivo counterparts, there are several key differences that are important to note. One such limitation is the gas environment of culture, which exposes cells to ambient oxygen levels (20%), much higher than normoxic levels of the brain (4-9%)^71,72^. Exposure to high oxygen can induce oxidative stress, requiring neuroprotective antioxidants to be added to cell culture media^73,74^. As such, cultures are likely exposed to an oxidative load that is close to the oxidative stress threshold and, thus, more sensitive to increases ROS production than in vivo tissues. We hypothesize that this high oxygen environment is the reason for the accelerated time course for oxidative damage in vitro compared to the longer in vivo disease progression. Additionally, exposure to high oxygen can have downstream effects on gene expression, which have yet to be characterized in such a model. Finally, it is important to acknowledge that while the absence of vascular-derived systemic immune cells in this model allows us to isolate and understand the innate immune response, it limits the scope of the conclusions that can be drawn about the in vivo microenvironment. Such in vitro testing helps to expand our understanding of disease states but does not preclude the necessity for in vivo experiments.

## Conclusion

The development and characterization of a novel 3D in vitro model of the device-tissue interface revealed a significant innate immune response to neural implants. This study has broad implications for the role of the innate immune system and potential treatments strategies for the chronic foreign body response to neural implants. Furthermore, the novel application of graph theory analysis to microcircuit calcium activity at the device-tissue interface exposed a breadth of underlying connectivity changes under neuroinflammatory conditions. The mitigation of oxidative stress, neuronal density, and functional connectivity with antioxidant ceria nanoparticles offers significant supporting evidence to indicate that many symptoms of the tissue reaction are related to redox dysregulation. Successful treatment of the device-tissue interface with long-acting antioxidant ceria nanoparticles shows potential for a local long-term antioxidant treatment strategy in clinical application, which could stabilize neural recordings and facilitate long-term use of implanted neural devices.

## Materials and Methods

### Animals

Animal handling and procedures were performed in accordance with Institutional Animal Care and Use Committee (IACUC) protocols, which were approved by Brown University and the Providence VA Medical Center. Sprague Dawley timed-pregnant rats sourced from Charles River Laboratories were monitored until pups were born. All biological experiments were performed using equal numbers of male and female rats. All procedures are in accordance with ARRIVE guidelines.

### Agarose Gels

Nonadherent microwell were made with 2% agarose (16500, Invitrogen) solutions using a custom injection molding system as previously described^50^. Briefly, a 2% agarose solution in phosphate buffered saline was heated in the microwave and mixed in 30 second intervals until fully dissolved. Hot agarose was then drawn into a 5mL syringe and injected though the custom mold and into the well of a glass bottom 24 well plate (P241.5HN, CellVis). Agarose was allowed to cool completely before removal of the injection mold. Gels were diffused with Complete Cortical Media (CCM) by incubation at 37C, with 2 media changes at a minimum interval of 2 hours.

### Three-dimensional Primary Cortical Cultures

Creation and maintenance of 3D primary cultured microtissues were performed as previously described^50^. In brief, primary cells were isolated from postnatal day 1 rats by cortical dissection in cold Hibernate A medium (HA, Brainbits) with B27 supplement (17504-044 Invitrogen) (HA-B27), followed by chemical dissociation with papain (PAP, Brainbits) in hibernate solution (HACA, Brainbits) at 30C for 30min with gentle inversion mixing every 5 minutes. A single cell suspension was made by trituration with a P1000 pipette in HA-B27. The cell suspension was filtered through a 40µm cell strainer and then centrifuged at 150rpm for 5 min. The HA-B27 supernatant was replaced with CCM, resuspending the cell pellet, and the filtering-centrifuging steps were repeated twice more. Live cells from an aliquot of the cell suspension were counted using a 1:1 mixture with Trypan Blue (T8154, Sigma-Aldrich), and then resuspended to a concentration of 2.7×10^8^ cells in 10uL of CCM. 10uL of suspended cells were seeded into each agarose microwell, and cells were allowed to settle in the incubator for 20 minutes before covering in 600uL CCM. Implanted microtissues were made by placing sterilized tungsten microwires in the bottom of the microwell and seeding overtop of the wire. Any adjustments to the wire position were made with sterile forceps immediately after seeding. Cultures were maintained in CCM at 37C 5%CO_2_ with 500uL media changes performed every other day starting at DIV1.

### AAV Transduction

AAV transduction of the genetically encoded GCaMP6s calcium reporter and mRuby control tag (AAV1-hSyn1-mRuby2-GSG-P2A-GCaMP6s-WPRE-pA, AddGene) was performed at DIV1 as previously described^50^. Briefly, replacement of CCM with 500uL viral media, made by diluting viral stock in CCM to a concentration of 1×10^6^vg/mL, was performed at DIV1 and allowed to incubate for 2 days before replacement with CCM on DIV4. A half CCM media change was performed on DIV5, followed by a full media change every other day for the remainder of the study.

### Lipid Peroxidation Assay and Analysis

Microtissues undergoing the lipid peroxidation assay (Image-iT Lipid Peroxidation Kit) were cultured until DIV 14. CCM was removed and replaced by 500uL of 10uM BODIPY fluorescent reagent in CCM and allowed to incubate for 30 minutes. After incubation, excess dye was removed and three quick washes with warm phosphate buffered saline were performed. Microtissues were immediately imaged in PBS at 590 (red) and 510 (green) emissions with an Olympus FV3000 confocal microscope. 10x image stacks of microtissues were collected and converted to TIF stacks in ImageJ. Analysis of whole-tissue and local lipid peroxidation was performed by creating a summed z projection of each channel and measuring the summed fluorescence of the projections within the whole-tissue or local ROI. The lipid peroxidation was determined by dividing the total measured fluorescence in the red channel by that of the green channel. Images in which the wire was not in the same z plane as the tissue due to improper wire placement or subsequent explantation were excluded from analysis. Additionally, malformed microtissues which did not form around the electrode were excluded from analysis.

### Live Image Acquisition

Live imaging of virally mediated fluorescence was performed with an Olympus FV3000 confocal microscope with a stage top environmental chamber attachment (WSKMX chamber, Tokai Hit) to keep cells at 37C with 5% CO2. Calcium and control tag fluorescence was measured at DIV 14, 16, and 18. GFP calcium fluorescence was recorded using a 10x objective with an open aperture at 15Hz. Laser power was adjusted each day to avoid overexposure. mRuby control fluorescent image stacks were collected with a 30x oil objective using a 1µm step size and optimized aperture. Laser power was adjusted to avoid overexposure.

### Calcium Image Processing and Analysis

Calcium videos were converted to tiff stacks in Image J using the Olympus Plugin, and maximum z projections of the videos were then saved as tiff images. Microtissues that were malformed or did not have tissue clearly in the same plane as the wire implant were excluded from analysis. Fluorescent cell bodies were identified as ROIs from z projections and masked videos were created from ROI coordinates as previously described^50^. Briefly, max projection images were smoothed using a Gaussian filter with a kernel size matching the cell body size in the 10x images. Binary images were created by adaptive thresholding and used to create a distance transform image. A Laplacian of Gaussian pyramid function was performed on the distance transform image to detect “blobs” or cell bodies. Manual interactive correction was performed to remove any cell bodies outside the tissue edge. For wire implanted microtissues, neural extensions which wrapped around the wire were frequently misidentified as cell bodies due to being obscured by the wire and were manually removed from ROI identification. ROI positions were used to make a binary mask, which was overlaid onto calcium videos. Calcium traces were extracted from masked videos using the FluoroSNNAP application^75^. DF/F traces created using a baseline fluorescent signal, which was identified as the 20^th^ percentile across a 10 second time window.

Graph theory analysis of calcium traces was performed as previously described^50^. In brief, time-series data of calcium fluorescence from each cell body in the image was used to create a cross correlation matrix in MATLAB with the corr() function. The cross-correlation matrix was then used to calculate the average node-pair correlation for each microtissue. The average clustering coefficient was calculated using a threshold-free weighted graph to measure the intensity of local connections, a method developed by Onnela et al^76^ and implemented in a MATLAB toolkit by Dingle et al^31^.The average pathlength of the weighted graph was determined by the shortest distance between nodes proportional to the network size, a method from Muldoon et al^77^ and implemented with a MATLAB toolkit by Dingle et al^31^. Modularity was determined by identifying modules through hierarchical clustering of node-pair correlations and calculating the sum of the correlation coefficients within a module divided by the sum of the expected correlation coefficients in a random network, a method developed by M.E.J. Newman^61^ and implemented with a MATLAB toolkit by Dingle et al^31^.

### Structural Image Processing and Analysis

3D 30x images were converted to tiff stacks in Image J using the Olympus Plugin. 3D images were binarized in Image J using automated Otsu thresholding. Summed z projections were then created and whole-tissue or local ROI’s were overlaid onto the image to measure the summed white voxels of the volume data. Volume fraction was calculated by dividing the measured summed white voxels by the total number of voxels. Microtissues that were malformed or did not have tissue clearly in the same plane as the wire implant were excluded from analysis. Several images from the oil objective acquisition were discovered to be blurred after collection due to an oil contamination issue and were excluded from analysis.

### Statistical Analysis

Statistical analysis was performed in GraphPad Prism. All biological data was collected from three independent replicate litters. All statistical tests performed were unpaired two-tailed t-tests. All data was tested for normality to determine the use of a parametric or non-parametric t-test. For all t-tests, each data point displayed represented a measured value from a single microtissue. All t-tests had a significance level of alpha<0.05 (*p<0.05, **p<0.01, ***p<0.001, ****p<0.0001), and all error bars represent the standard deviation from the mean.

## Data Availability

All data generated or analyzed during this study will be made publicly available upon publication and can be made available upon request.

## Acknowledgements

Geoff Williams for his guidance and technical support with image acquisition, Diane Hoffman-Kim and Liane Livi for their guidance and support with cell culture, Ben Goddard for his guidance and technical support with data analysis.

## Author Contributions

E.A., V.C. and D.A.B. conceived of the experiments; E.A., Y.H., S.B., E.P. and V.L. performed the experiments; E.A. and S.B. performed image acquisition and data analysis; E.P. and E.A. performed the toxicity tests; Y.H. synthesized the ceria formulations; Y.H. and V.L. performed the ceria characterization; E.A. wrote the first draft of the manuscript; All authors reviewed and edited the final manuscript.

## Competing Interests

The authors declare no competing interests.

## Funding

This work was supported in part by the Center for Neurorestoration and Neurotechnology (N2864-C) from the United States (U.S.) Department of Veterans Affairs, Rehabilitation Research and Development Service, Providence, RI. The contents do not represent the views of the U.S. Department of Veterans Affairs or the United States Government.

## Notes

### Competing Interest Statement

The authors have declared no competing interest.

