## Supplemental Figures for "Three-dimensional in vitro model of the device-tissue interface reveals innate neuroinflammation can be mitigated by antioxidant ceria nanoparticles"

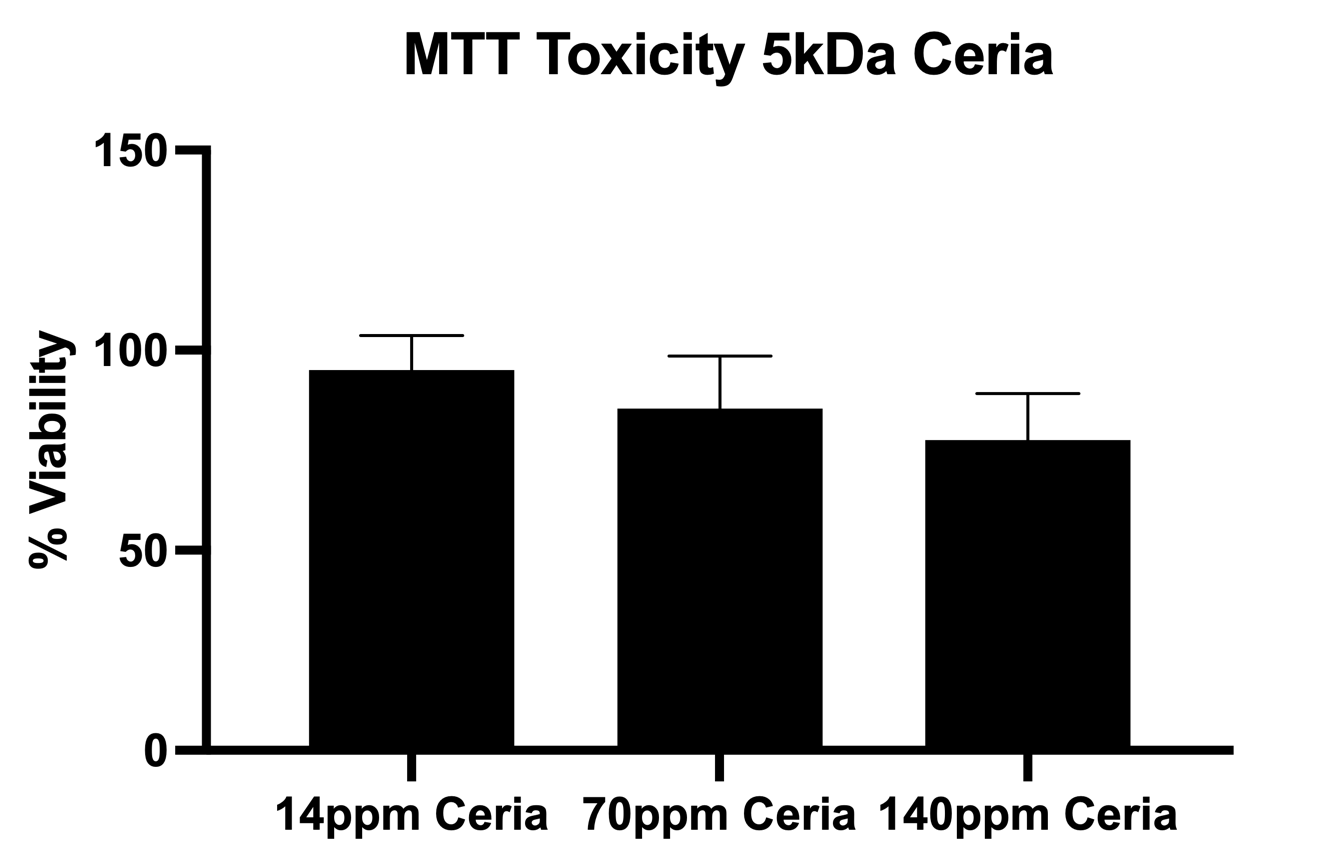

**Figure S1. Ceria nanoparticle toxicity.** MTT assays, which measure mitochondrial function, performed at 24 hours after a single concentration exposure of 14ppm, 70ppm, or 140ppm ceria show that 14 ppm has the lowest effect on normal mitochondrial function.

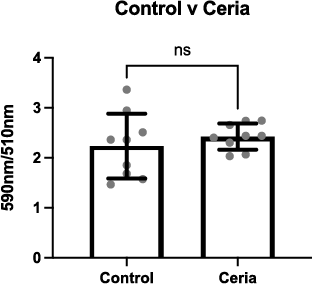

**Figure S2. Oxidative stress in ceria-treated unimplanted microtissues.** Ceria treatment of unimplanted microtissues showed no significant difference in lipid peroxidation (p = 0.4341) compared to unimplanted microtissues that did not receive ceria.

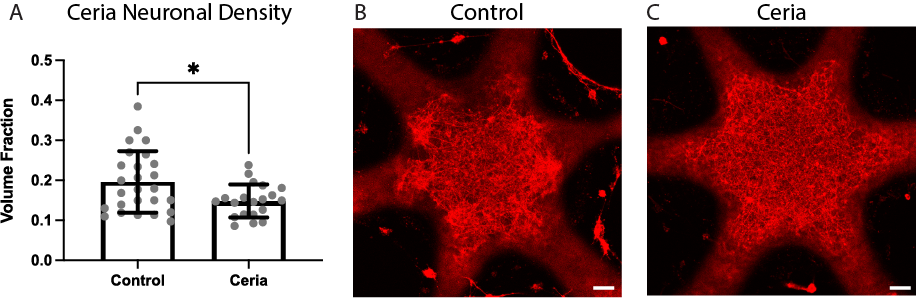

**Figure S3. Ceria treatment of unimplanted microtissues causes neuronal density changes.** (A)Ceria treatment in uninjured tissues significantly decreases the neuronal density (p=0.0163). Structural neuronal changes can be seen visually in (B) controls compared to (C) ceria treated microtissues, which show more regular distribution of neuronal extensions. Scale bars = 100µm; *p<0.05.

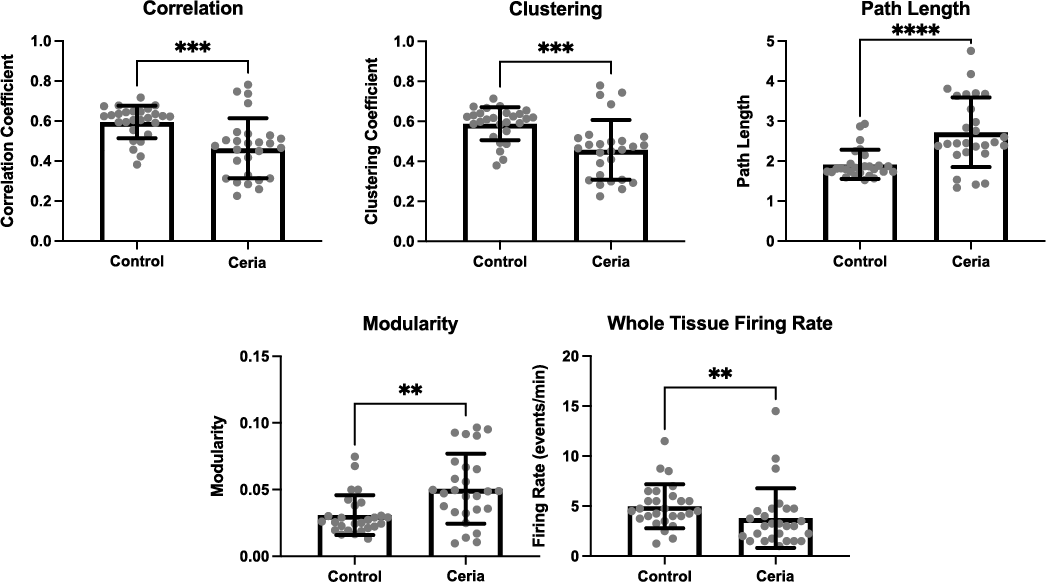

**Figure S4. Whole-tissue ceria treatment of unimplanted microtissues causes functional connectivity and community structure changes.** Ceria treatment of unimplanted microtissues compared to untreated controls showed a significant decrease in (A) Correlation (p=0.0002) and (B) Clustering(p=0.0002), and a significant decrease in (C) Path Length (p<0.0001). Ceria treatment also increased (D) Modularity (p=0.0013) and decreased the (E) Whole-tissue firing rate(p=0.0058). **p<0.01, ***p<0.001, ****p<0.0001.

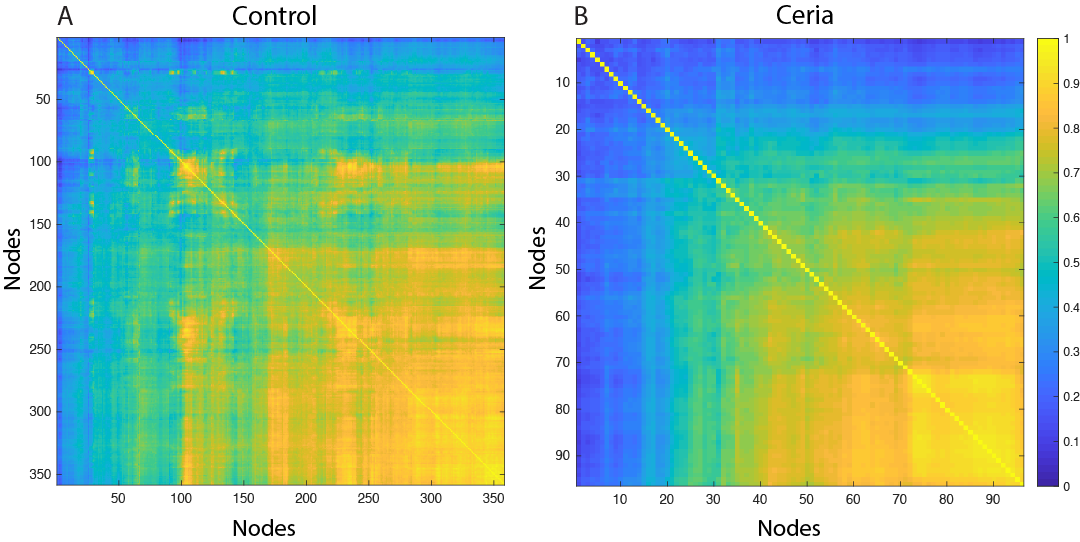

**Figure S5. Decreased functional connectivity in ceria treated microtissues.** Correlograms of representative (A) control and (B) ceria treated microtissues show the decreased in correlation induced by ceria treatment in unimplanted microtissues.
